# RNA exosome ribonuclease DIS3 degrades *Pou6f1* to promote mouse pre-implantation cell differentiation

**DOI:** 10.1101/2022.05.01.490217

**Authors:** Di Wu, Jurrien Dean

## Abstract

Mammalian development is precisely controlled by cell differentiation. Discovery of new regulators and investigating their crosstalk provide insights into genetic networks defining pre-implantation development. To identify novel developmental repressors, we established a knockout mouse model of *Dis3*, an RNA exosome associated RNase. Homozygous *Dis3* null embryos arrest at the morula-to-blastocyst transition. Using single embryo RNA-seq, we discovered persistence of *Pou6f1* mRNA in homozygous null *Dis3* embryos and determined that the cognate protein represses transcription of *Nanog* and *Cdx2*. The resultant defects in cell differentiation disrupted the morula-to-blastocyst transition and was embryonic lethal. To overcome the paucity of embryos, homozygous *Dis3* null mouse embryonic stem cells were derived to identify additional gene targets of POU6F1. While microinjection of the *Dis3* cRNA into zygotes rescued the morula-to-blastocyst block, point mutations of *Dis3* in individual blastomeres transformed the cell and prevented incorporation into embryos. Our findings uncover a not heretofore reported regulatory pathway of DIS3-POU6F1 in pre-implantation mammalian embryogenesis.

**In Brief:** Mammalian pre-implantation development is regulated by master transcription factors and their crosstalk. Wu and Dean report that an RNA exosome associated RNase, DIS3, degrades *Pou6f1* mRNA to de-repress transcription of *Nanog* and *Cdx2* genes. In the absence of DIS3, POU6F1 protein persists and embryos arrest as morulae unable to become blastocysts due to lack of cell differentiation.

**Highlights:**

- *Dis3* knockout mice have morula arrest due to lack of cell differentiation.
- DIS3 binds and degrades *Pou6f1* mRNA before the morula stage.
- POU6F1 globally occupies promoters to regulate gene transcription.
- DIS3 mutation results in cell transformation in embryonic development.

## INTRODUCTION

The maternal-to-zygotic transition in early mammalian development is conserved and includes maternal RNA degradation as well as zygotic gene activation (ZGA) (Jukam et al., 2017; Svoboda et al., 2015). Failure to clear maternal RNAs results in embryonic arrest (Sha et al., 2020) and perturbation of ZGA prevents progression beyond 2-cell embryos. Loss-of-function studies document critical licensing factors that control zygotic gene transcription (Svoboda et al., 2015), but less is known about the molecular regulation governing clearance of maternal RNA.

The RNA exosome complex is responsible for degrading RNA in many different cell types. The well-conserved exosome complex binds to associated RNases (DIS3, EXOSC10, DIS3L) to acquire enzymatic activity (Kilchert et al., 2016; Zinder and Lima, 2017). One of the most important RNA exosome-associated RNase, DIS3, is essential for cell survival and embryo development in flies (Hou et al., 2012). Depletion of DIS3 causes accumulation of a wide range of RNAs, including protein coding RNA, noncoding RNA, and repetitive element RNA. Using CLIP-seq, the direct RNA targets of DIS3 have been reported in human and mouse cultured cell lines (Davidson et al., 2019; Szczepinska et al., 2015). Although target selection of DIS3 ribonuclease lacks sequence specificity, the binding profiles exhibit strong tissue preference because of distinct transcriptomes in different cells (Davidson et al., 2019; Hou et al., 2012; Szczepinska et al., 2015; Wu and Dean, 2022). Thus, preventing RNA degradation by DIS3 ribonuclease depletion can identify regulators of development when their persistence causes significant downstream effects.

During mammalian pre-implantation development, the embryo undergoes zygotic gene activation, cell cleavage, compaction, cavitation, and differentiation (Wu and Dean, 2020). At the end of the blastocyst stage, three cell lineages are defined by position and expression of unique markers, namely EPI (epiblast, expressing NANOG), PrE (primitive endoderm, expressing GATA4/GATA6), and TE (trophectoderm, expressing CDX2/GATA3) (Schrode et al., 2013). Master transcription factors, including SOX2, OCT4, NANOG, and CDX2 (Avilion et al., 2003; Niwa et al., 2005; Ralston and Rossant, 2008), drive serial differentiation during two sequential cell fate decisions, namely the ICM (inner cell mass) versus the TE (trophectoderm) differentiation and EPI/PrE delineation within the ICM (Schrode et al., 2013). The mutual activation and repression between different factors synchronize different cell lineages for future germ layers. For example, OCT4 and SOX2 directly initiate *Nanog* transcription (Rodda et al., 2005), and *Nanog* and *Cdx2* mutually repress each other’s transcription by occupying promoter regions (Chen et al., 2009). Although the molecular mechanism underlying the dynamic sculpting of different cell fates has been explored extensively, the complexity remains incompletely understood.

In this study, we constructed *Dis3* knockout mice and homozygous embryos arrest during pre-implantation development. We demonstrate that the defects occur in cell differentiation during the morula-to-blastocyst transition. Through single embryo RNA-seq analysis, we identified POU6F1, a transcription factor, as a potential target for DIS3 mediated degradation. We profiled gene targets of POU6F1 to confirm its repressive activity on transcription. Finally, we tested the effects of mutant forms of DIS3 ribonuclease which provide insights into formation of congenital developmental abnormalities and oncogenesis.

## RESULTS

### *Dis3* Homozygous Null Embryos Arrest at the Morula-to-Blastocyst Transition

A *Dis3* null allele was fortuitously generated while establishing mouse lines with a floxed allele. Exons 3-5 were deleted by CRISPR/Cas9 and imprecisely repaired through non-homologous end joining (Figure 1A, S1A, S3C) to create the null allele. We crossed heterozygous *Dis3* mice (het) and none of the pups born were homozygous null *Dis3* (null), suggesting embryonic lethality (Figure 1B). In contrast, *Dis3* het mice were healthy (Figures 1B, S1B) and grew to adulthood.

**Figure 1.**
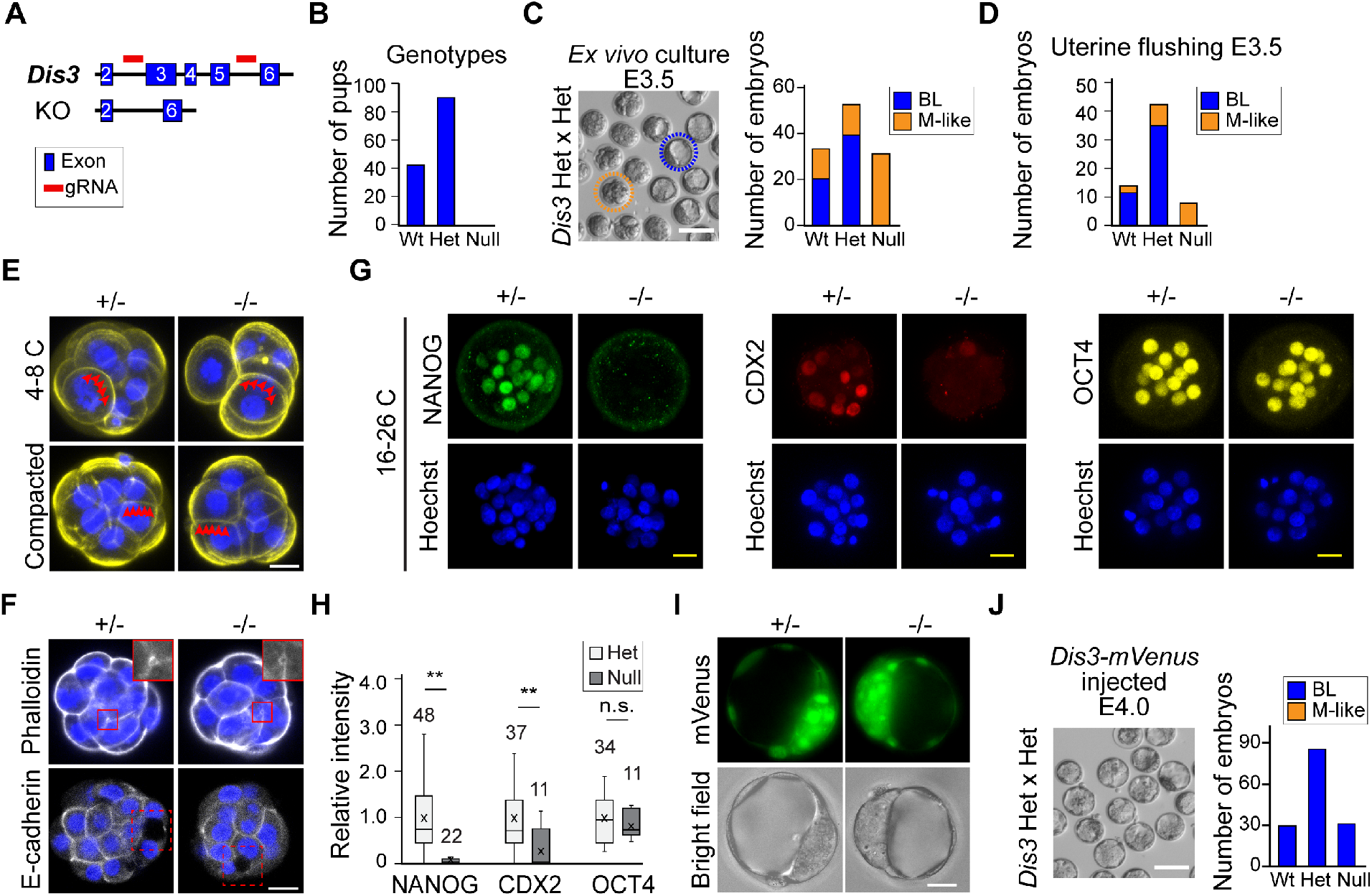
*Dis3* Null Embryos Arrest at the Morula-to-blastocyst Transition and Lack Cell Differentiation. (A) Schematic of *Dis3* knockout mice construction. Exons 3-5 are absent in the *Dis3* null allele. (B) Bar graph showing the number of pups of each genotype born from *Dis3* heterozygous mating. (C and D) Number of blastocysts (BL) and morula-like embryos (M-like) in *ex vivo* culture (C) and uterine flushing (D) at E3.5 derived from *Dis3* heterozygous mating. (E) Phalloidin staining of *Dis3* heterozygous (+/-) and null (-/-) embryos at the 4-8 cell stage before and after compaction. Red triangles indicate boundaries of blastomeres. (F) Immunofluorescence of het (+/-) and null (-/-) embryos after staining with phalloidin and E-cadherin. Insets, regions enlarged to show the microlumen. (G and H) Immunofluorescence and quantification of NANOG, CDX2 and OCT4 in *Dis3* het (+/-) and null (-/-) embryos. ** NANOG, P=2.2E-10; ** P=CDX2, 0.0006; n.s., not significant; two-tailed Student’s t test. (I) mVenus fluorescence of the injected *Dis3-mVenus* cRNA in *Dis3* het (+/-) and null (-/-) embryos. (J) Number of BL and M-like embryos in *ex vivo* culture at E3.5 of embryos derived from *Dis3* heterozygous mating and microinjected with *Dis3-mVenus* cRNA at 1-cell stage. Scale bars: 100 μm in C, J; 20 μm in E, F, G, I.

We sought to identify the stage of lethality of *Dis3* null embryos. In *ex vivo* culture of the pre-implantation embryos produced by *Dis3* heterozygous mating, we observed a decrease of viability at embryonic day 4 (E4) (Figure S1C). Around 25% of the embryos arrested at the morula stage and could not form a blastocoel. We defined the embryos as either blastocyst (BL, having an obvious blastocoel) or morula-like (M, not having an obvious blastocoel) at E3.5, and performed single embryo genotyping. All *Dis3* null embryos belonged to the morula-like group (Figure 1C). The same morula-arrest phenotype could be recapitulated by uterine flushing of female mice at E3.5 after *Dis3* heterozygous matings (Figure 1D). To further validate the arrest, we synthesized *Dis3* morpholino (MO2) and microinjected it into 1-cell wildtype embryos to block translation of *Dis3* mRNA (Figures S1D-E). As expected, *Dis3* morpholino efficiently blocked the translation of its downstream reporter gene, and *Dis3* morpholino injection increased morula-like embryos and reduced blastocysts at E4.0 in a dose-dependent manner (Figures S1F-H).

To further investigate the cause of the morula arrest, we confirmed that cells did not undergo apoptosis in *Dis3* null embryos at E3.5 (Figures S2A-B). By outlining the cell shape by phalloidin staining, we observed successful compaction in *Dis3* null embryos (Figure 1E). The *Dis3* null embryos also initiated blastocoel cavitation by forming micro-lumens visualized by phalloidin and E-cadherin immunostaining (Figure 1F). In addition, *Dis3* null embryos did not have obvious differences in their endomembrane systems (Figures S2C). However, when we performed immunostaining of NANOG and CDX2, which are cell differentiation markers, *Dis3* null embryos had significantly reduced NANOG and CDX2 (Figures 1G-H, S2D) which was not a reflection of embryonic stage difference. Another differentiation marker, OCT4, remained normal in *Dis3* null embryos due to its maternal origin (Figures 1G-H). To further document that DIS3 was essential for the morula-to-blastocyst transition, we synthesized *Dis3-mVenus* cRNA *in vitro* and microinjected it into the 1-cell embryos derived from *Dis3* heterozygous mating. Overexpression of *Dis3-mVenus* cRNA resulted in an increase in the number of blastocysts because the *Dis3* null embryos successfully formed a blastocoel (Figure 1I-J).

### DIS3 Degrades *Pou6f1* mRNA to De-repress *Nanog* and *Cdx2* Transcription

The reduction of NANOG and CDX2 suggested an inhibition of their gene transcription in *Dis3* null embryos. We hypothesized that depletion of DIS3 ribonuclease results in insufficient degradation of mRNA encoding a repressor which consequently blocked gene transcription prior to the morula stage. To profile potential DIS3 target genes, we performed single embryo RNA-seq at E2.5 and E2.75. As expected, *Dis3* het and wildtype embryos were grouped together by PCA (principal component analysis), whereas *Dis3* null embryos at both stages were in a distinct group (Figure S3A). Due to the similarity of the differential genes at the two stages, we combined them for a differential analysis. As a result, we obtained 1,245 up-regulated genes and 547 down-regulated genes in null vs. het comparison, and only 37 up-regulated and 29 down-regulated genes in het vs. wildtype comparison (Figure 2A). The decrease of *Nanog* and *Cdx2* transcripts was also apparent in the differential analyses (Table S2). In addition to the coding RNAs, PROMPTs RNA, which are known DIS3 targets, also exhibited significant accumulation (Figure S3B) (Davidson et al., 2019; Szczepinska et al., 2015).

**Figure 2.**
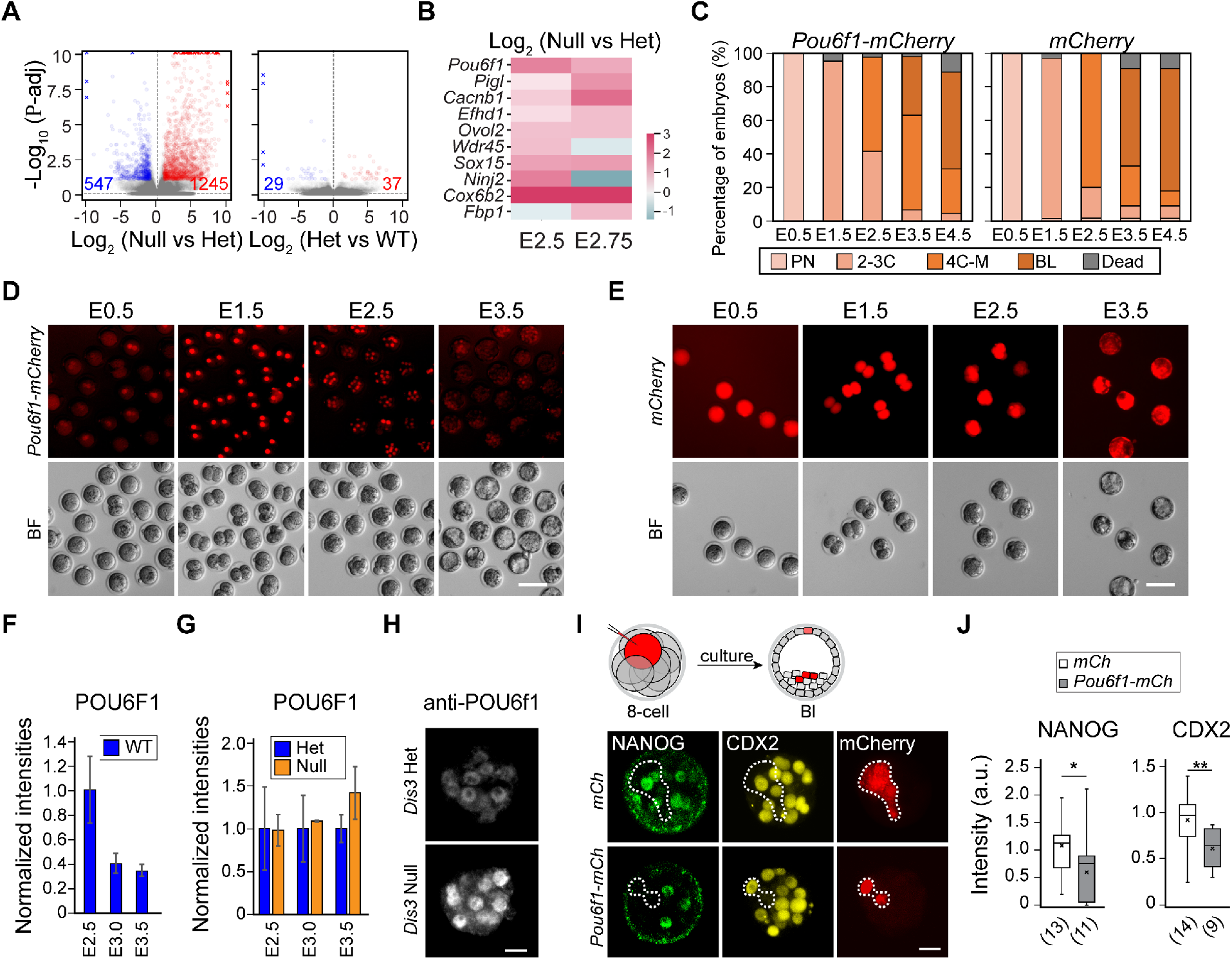
*Pou6f1* is Up-regulated in *Dis3* Null Embryos and Blocks *Nanog* and *Cdx2* Transcription. (A) Volcano plots showing differentially expressed genes from single embryo RNA-seq of combined analyses of E2.5 and E2.75. Left, *Dis3* null embryos vs heterozygotes (Null vs Het); right, *Dis3* heterozygotes vs wildtype (Het vs WT). Blue and red, significantly down- and up-regulated genes (P-adj <0.1). (B) Heatmap showing the change of 10 selected genes in Null vs Het at E2.5 and E2.75. (C) Bar graph showing the ratio of embryos at each stage from E0.5 to E4.5 in *ex vivo* culture. Wildtype 1-cell embryos (around 20) were injected either with *Pou6f1-mCherry* (N=21) or *mCherry* (N=19). Results combine two assays. (D and E) Florescent and bright field images of embryos from E0.5 to E3.5 that were either *Pou6f1-mCherry* injected or *mCherry* injected. Scale bars: 100 μm. (F) Quantification of POU6F1 immunofluorescence staining in WT embryos from E2.5 to E3.5. (G and H) Quantification and immunofluorescence staining of POU6F1 in *Dis3* het and null embryos (normalized by het embryos). (I and J) Schematic, immunofluorescence staining and quantification of NANOG and CDX2 in blastomeres injected with *Pou6f1-mCherry* or *mCherry* (in mCherry positive cells). * NANOG, P=0.096; ** CDX2, P=0.009. Scale bars: 20 μm in H, I.

To screen for DIS3 substrates, we took advantage of published DIS3 CLIP-seq datasets from cultured cell lines to overlap with the protein-coding genes that were significantly increased in at least one stage in our RNA-seq results (Figures 2B, S3D) (Szczepinska et al., 2015). From the obtained gene lists, we *in vitro* synthesized 10 cRNAs as mCherry fusion genes, and overexpressed the cRNAs individually by microinjection into 1-cell wildtype embryos (Figures 2C-E, S4A). We observed that *Pou6f1* overexpression increased morula arrest at E3.5 and consequently there were fewer blastocysts at E4.5 (Figures 2C-E). From public datasets (GSE57249 and GSE45719), we confirmed that RNA levels of *Pou6f1* increased in wildtype embryos as they advanced from 1-to 4-cell stage and then decreased in the blastocyst stage (Biase et al., 2014; Deng et al., 2014) with the cognate protein decreasing from E2.5 to E3.0 (Figures S3E-G), suggesting the need for POU6F1 clearance before the blastocyst stage. Degradation of *Pou6f1* transcripts was prevented in *Dis3* null embryos, which resulted in persistence of POU6F1 protein (Figures 2B, F-H).

To validate the inhibitory effect of POU6F1 on *Nanog* and *Cdx2* transcription, we microinjected *Pou6f1-mCherry* cRNA into 1-2 blastomeres at the 8-cell stage and measured NANOG/CDX2 at the blastocyst stage by immunostaining (Figure 2I). As expected, cells injected with *Pou6f1-mCherry* cRNA had lower NANOG and CDX2 intensities compared to uninjected cells in the same embryo (Figure 2J). We conclude that *Dis3* null embryos fail to degrade *Pou6f1* which consequently repressed the transcription of *Nanog* and *Cdx2* genes.

### POU6F1 Binds and Represses Gene Transcription in Mouse Embryonic Stem Cells

Our results suggested possible roles of DIS3 and POU6F1 in regulating cell differentiation. To directly test this hypothesis, we derived mouse embryonic stem cells (mESCs) from the *Dis3^Flox/Flox^* blastocysts (Figure 3A). The obtained mESCs were positive for stem cell markers OCT4, NANOG and SSEA-1, and negative for the differentiation marker, NESTIN (Figure 3B). mESCs were able to maintain their stemness in self-renewal medium and underwent differentiation upon removal of self-renewal factors (Figures S5A-B). The differentiated mESCs became negative for NANOG and positive for NESTIN as well as the neural ectodermal marker PAX6 (Figure S5C). When DIS3 was depleted by transfecting cre recombinase with the CAG promoter into the mESCs, we observed a significant upregulation of *Pou6f1* when cells were in differentiated state (Figure 3C). To validate the ability of DIS3 to bind *Pou6f1* mRNA, we synthesized the DIS3-occupied fragment of *Pou6f1* from the published CLIP-seq dataset to perform electrophoretic mobility shift assay (EMSA) (Szczepinska et al., 2015) (Table S1, Figure S3D). A mutant form of DIS3 protein was used to avoid degradation after binding in the EMSA (DIS3-DN-ΔNLS-mVenus, Figures 3D, 4A-B). As a result, DIS3 protein bound to the synthesized *Pou6f1* fragment and was super-shifted after addition of anti-FLAG antibody (Figure 3D) to validate the interaction.

**Figure 3.**
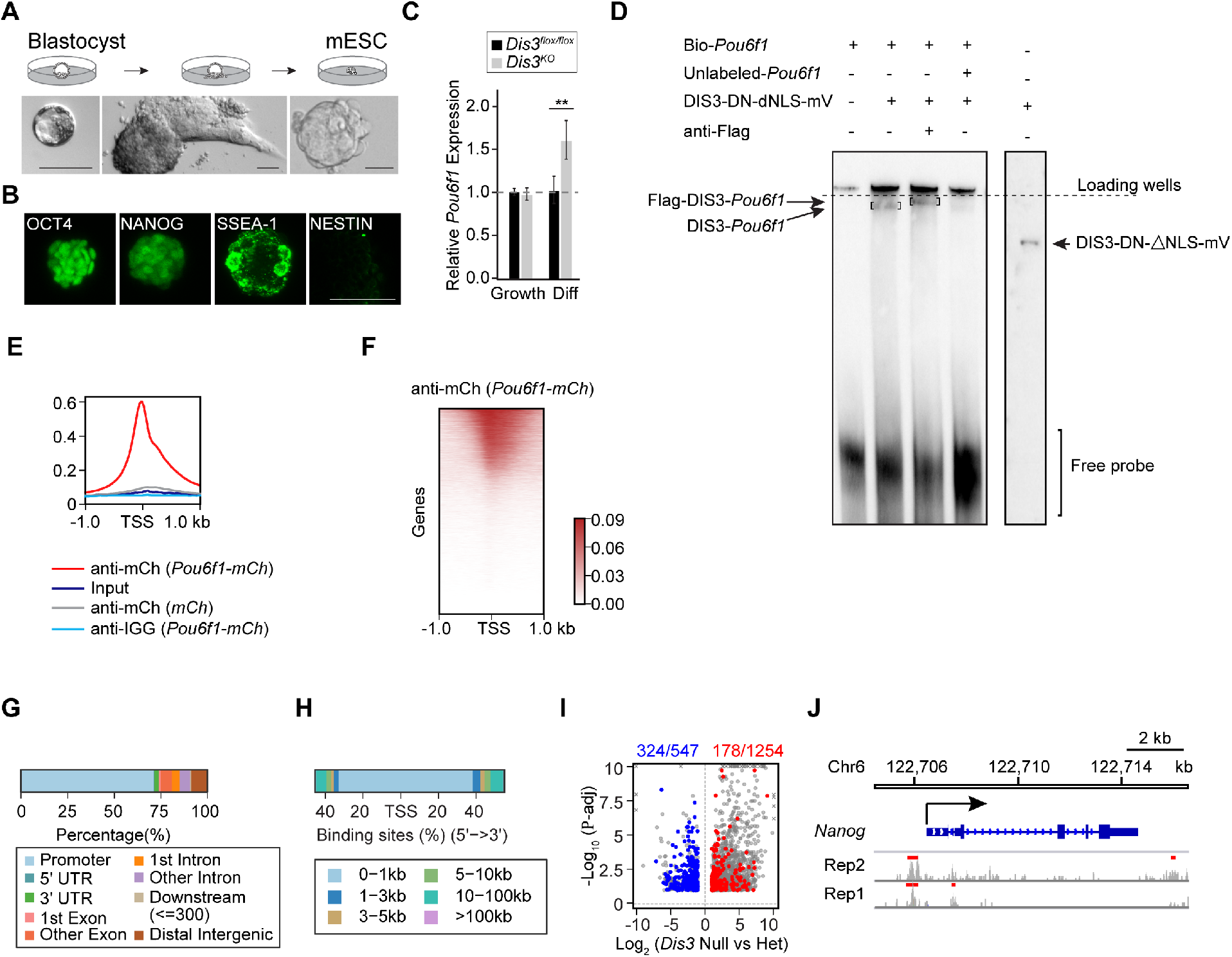
POU6F1 Globally Occupies Promoters to Repress Gene Transcription in Mouse Embryonic Stem Cells. (A) Schematic of mouse embryonic stem cells (mESC) derivation from *Dis3^flox/flox^* blastocysts. (B) Immunofluorescence staining of OCT4, NANOG, SSEA-1 and NESTIN in mESCs. (C) Bar graph showing *Pou6f1* changes in growth and differentiation (Diff) phases of mESCs after knockout of *Dis3*. **, P=0.01. (D) Electrophoretic Mobility Shift Assay (EMSA) result showing DIS3-DN-ΔNLS-mVenus (D146N+D487N) protein binding to *Pou6f1* RNA. On the right shows the protein band detected in western blot using anti-Flag (engineered epitope in the mVenus protein). (E and F) Profile and heatmap of POU6F1 occupancy from ChIP-seq analysis. (G and H) Distribution of POU6F1 occupied genome loci in different genomic regions and the relative distances of the peaks to TSS. (I) Volcano plot of significantly changed genes from RNA-seq (from Figure 2A) having POU6F1 occupancy. Blue and red, significantly changed down- and up-regulated genes in *Dis3* null vs heterozygotes that are occupied by POU6F1. (J) POU6F1 peaks on *Nanog*. Rep1 and 2: two biological replicates. Red bars: peaks called against anti-IGG samples. Scale bars: 100 μm in A; 50 μm in B.

**Figure 4.**
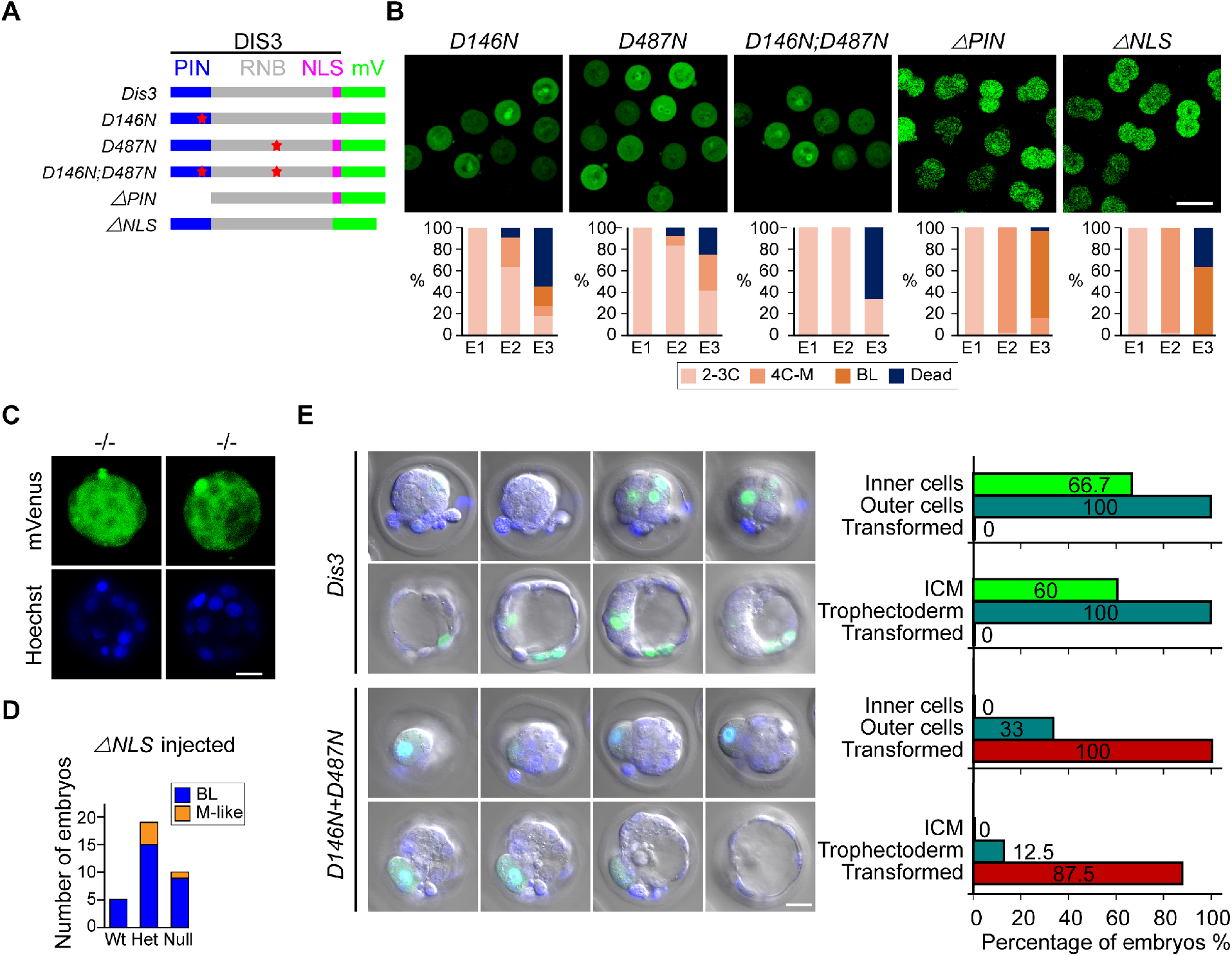
Mutant Forms of DIS3 Can Induce Cell Transformation in Early Embryonic Development. (A) Schematic of all DIS3 mutant forms of protein. PIN, DIS3 PIN domain; RNB: DIS3 RNB domain; NLS: DIS3 nuclear localization signal; mV: mVenus coding region. (B) Fluorescent live imaging of DIS3 mutant proteins and the ratio of embryos at different stages from E1 to E3. (C) Fluorescent live imaging of DIS3-ΔNLS-mV protein at the blastocyst stage in Dis3 *null* embryos. Embryos were microinjected with *Dis3-ΔNLS-mV* cRNA, cultured, and imaged individually and collected for single embryo genotyping. (D) Ratio of BL and M-like embryos from *Dis3* heterozygous mating and microinjected with *Dis3-ΔNLS-mV* cRNA. (E) Fluorescent live imaging and cell position quantification after microinjected with *Dis3-mV* or *Dis3-D146ND487N-mV* cRNA into 1-2 blastomeres at the 4-8 cell stage. Cell positions were visually identified as inner, outer, and transformed based on their position and size. Scale bars: 100 μm in B; 20 μm in C, E.

Next, we performed ChIP-seq of POU6F1 in mESC cells to profile target genes. Two biological replicates exhibiting high reproducibility were combined for analysis of POU6F1 binding sites (Figure S5D-E) (Table S3). Globally, we have identified around 18,000 peaks (Table S3). POU6F1 binding sites were close to transcription start sites (TSS) (Figures 3E-F). More than 70% of the POU6F1 peaks were present in promoter regions (Figure 3G) and more than 80% peaks were within 1 kb of the TSS (Figure 3H). When overlapping the POU6F1-occupied genes with the differential genes from the *Dis3* embryonic RNA-seq, ~60% (324/547) of the down-regulated genes were occupied by POU6F1, while only ~15% (178/1,254) of the up-regulated genes were occupied by POU6F1 (Figure 3I). From the POU6F1 occupied promoters, there were 317/9119 genes involved in *in utero* embryonic development (q-value: 1.5E-26, Table S4). Specifically, we observed occupancy of the repressive transcription factor POU6F1 in *Nanog* and *Cdx2* promoters and genic regions (Figures 3J, S5F). To summarize, *Pou6f1* mRNA is degraded by DIS3 and its persistence in the absence of DIS3 ribonuclease is associated with continued repression of differentiation genes with subsequent embryonic arrest.

We were then interested in exploring potential interaction proteins of POU6F1. From two public databases, namely BioGRID (Oughtred et al., 2021) and IntAct (Hermjakob et al., 2004), we found three interacting partners of POU6F1, including ACOT7, ERBB2 and NFYC. Among them, ERBB2 and NFYC are nuclear proteins and have known target genes (Fleming et al., 2013; Redmond et al., 2019). We overlapped the ERBB2 and NFYC occupied genes with POU6F1 occupied genes, respectively. About 27% (2,516/9,302) of POU6F1 occupied genes were overlapped with ERBB2, about 43% (3,997/9,302) of POU6F1 occupied genes were overlapped with NFYC, and 1,602 genes were shared by the three genes (Figure S5G). The overlap in occupancy suggests a potential cooperation of POU6F1 with the other factors on transcriptome regulation.

### Mutant DIS3 Proteins Induce Cell Transformation

Mutations of DIS3 have been reported in several human diseases, including multiple myeloma and acute myeloid leukemia (Ding et al., 2012; Weissbach et al., 2015). We engineered the wildtype DIS3 protein into several mutant forms, including an endonucleolytic mutation (D146N), an exonucleolytic mutation (D487N), a double mutation (D146N+D487N), an endonucleolytic deletion (ΔPIN) and a nuclear localization signal deletion (ΔNLS) (Figure 4A). After microinjection into 1-cell wildtype embryos, D146N, D487N, and D146N+D487N caused embryonic arrest at the 2-cell stage (Figure 4B). However, ΔPIN and ΔNLS overexpression did not affect pre-implantation development (Figure 4B). Interestingly, DIS3-ΔNLS overexpression could rescue *Dis3* null embryos to form blastocysts which further supports a model in which accumulated cytoplasmic RNA, instead of nuclear RNA (including pervasive transcripts), was more likely to cause the morula arrest (Figures 4C-D).

To determine the effect on single blastomeres in embryos, we microinjected wildtype *Dis3* or *D146N+D487N* cRNA into 1-2 blastomeres at the 8-cell stage and recorded the position and morphology of the injected blastomeres at later stages. Blastomeres injected with wildtype *Dis3* cRNA were present in either inner or outer positions in morula and blastocysts (Figure 4E). In contrast, blastomeres injected with *D146N+D487N* cRNA were all restricted to the embryo surface (Figure 4E). They did not appear as “outer cells” because their sizes were significantly larger than uninjected cells in the same embryo. Instead, we defined them as transformed cells due to their inability to be incorporated into dividing embryos, which may be due to dramatic transcriptome disruption and/or concurrent genomic instability (Milbury et al., 2019). We conclude that expression of mutant D146N+D487N directly results in embryonic arrest or cell transformation in pre-implantation development.

## DISCUSSION

Our results demonstrate the essential role of RNA degradation in early embryo development. Deletion of *Dis3*, an RNA exosome associated RNase, results in developmental arrest at the morula-to-blastocyst transition. This led to the identity of *Pou6f1* mRNA as an important target of DIS3 ribonuclease at this stage of development. The persistence of *Pou6f1* mRNA due to insufficient degradation leads to more abundant POU6F1 protein that represses transcription of *Nanog* and *Cdx2*, and blocks cell differentiation

Mutual cooperative interactions and/or antagonistic regulation of master transcription factors is a hallmark of sequential cell fate specification that defines cell lineage. A transcription regulator can not only prime cells for downstream differentiation but also repress their proclivity towards other cell fates. For example, OCT4 (POU5F1) activates *Nanog* transcription in future ICM cells and represses *Cdx2* transcription to prevent a PE fate. In contrast, in future PE cells *Oct4* and *Nanog* are repressed by CDX2. In the current manuscript, we identify POU6F1, another member of the POU-homeobox domain family transcription factors, as actively involved in this regulatory network. POU6F1 represses both *Nanog* and *Cdx2* gene expression at the morula stage in wildtype embryos which suppresses stemness and induces cell differentiation. If *Pou6f1* mRNA is not degraded, cells cannot differentiate and embryos arrest as morulae during pre-implantation development. Similar observations were made in mESCs cultured *in vitro*. Consistent with these results, POU6F1 has also been reported essential for specification and patterning of dendrites, and its overexpression can promote dendritic outgrowth and branching (McClard et al., 2018).

Our findings demonstrate that zygotic DIS3 is essential for the morula-to-blastocyst transition during pre-implantation development. The level of *Dis3* transcript is decreased in *Dis3* null embryos from E2.5 onward. However, it does not rule out a function for DIS3 before E2.5 due to maternal persistence of *Dis3* RNA and protein. To address this issue, we synthesized a *Dis3* MO to block maternal RNA translation which could expose an earlier phenotype by reducing maternal DIS3 protein level. However, we obtained the same morula arrest phenotype in *Dis3* MO injected embryos which suggested that maternal *Dis3* RNA is not essential for early embryogenesis. Whether maternal DIS3 protein regulates pre-implantation development remains unknown, because conditional depletion in oocyte of DIS3 blocks maturation and precludes studying a post-fertilization maternal effect (Wu and Dean, 2022).

The RNA exosome complex is responsible for degrading many types of RNA. Multiple studies have focused on finding a sequence-based affinity of DIS3-bound RNAs without success, suggesting that the recognition and degradation by DIS3 does not depend on sequence defined domains (Davidson et al., 2019; Szczepinska et al., 2015). We reasoned that one or more persistent transcripts could cause a particular phenotype but only under certain cellular conditions or in combination with other perturbations. Indeed, among the 10 genes we screened from the upregulated transcripts in *Dis3* null embryos, only *Pou6f1* overexpression partially phenocopied the morula arrest. Due to the paucity of biological material, we profiled POU6F1 genomic occupancy in mESCs rather than in morula embryos. Nevertheless, we identified many POU6F1 occupied promoters, whose genes are involved in cell differentiation and embryogenesis, which strongly supports that a model in which POU6F1 binds to chromatin and regulates cell fate transition. Based on our results with mutant forms of *Dis3*, it appears that cytoplasmic DIS3 can rescue blastocyst formation which excludes a role for nuclear RNAs in triggering morula arrest, and further supports a critical cytoplasmic function of DIS3 in degrading mRNAs.

DIS3 mutations and altered protein level have been reported in several human diseases (Ding et al., 2012; Milbury et al., 2019; Weissbach et al., 2015). One of the known enzymatic sites of DIS3 RNB domain (D487) was tested in early embryonic development by overexpression and led to 2-cell arrest. In addition, DIS3 with mutations in both endonuclease and exonuclease domains (D146N, D487N) caused cell transformation and exclusion from embryos. This may provide a possible mechanism of DIS3 mutations causing genomic instability and genome rearrangement in human diseases. Interestingly, dysregulated genes in *Dis3* null embryos include *MTAP, G3BP2, SEC23IP* and *USO1*, whose changes were similar to superficial spreading melanoma, the most common form of melanoma (GSE22301) (Rose et al., 2011), which has decreased *Dis3* expression. Our study not only elucidates the essential role of DIS3 in early embryonic development but also highlights its conserved roles across different cellular contexts in maintaining transcriptome integrity.

Collectively, our results demonstrate the essential role of exosome DIS3 ribonuclease in mammalian early embryogenesis. We identified *Pou6f1* mRNA as a target of DIS3 that transcriptionally represses genes necessary to establish cell lineages. Insufficient degradation of *Pou6f1* leads to loss of cell differentiation and embryonic arrest. In summary, these results incorporate a not heretofore reported regulatory role for DIS3 in regulating the abundance of POU6F1 to facilitate pre-implantation development in mammals.

## STAR METHODS

Detailed methods are provided in the online version of this paper and include the following:

### KEY RESOURCES TABLE

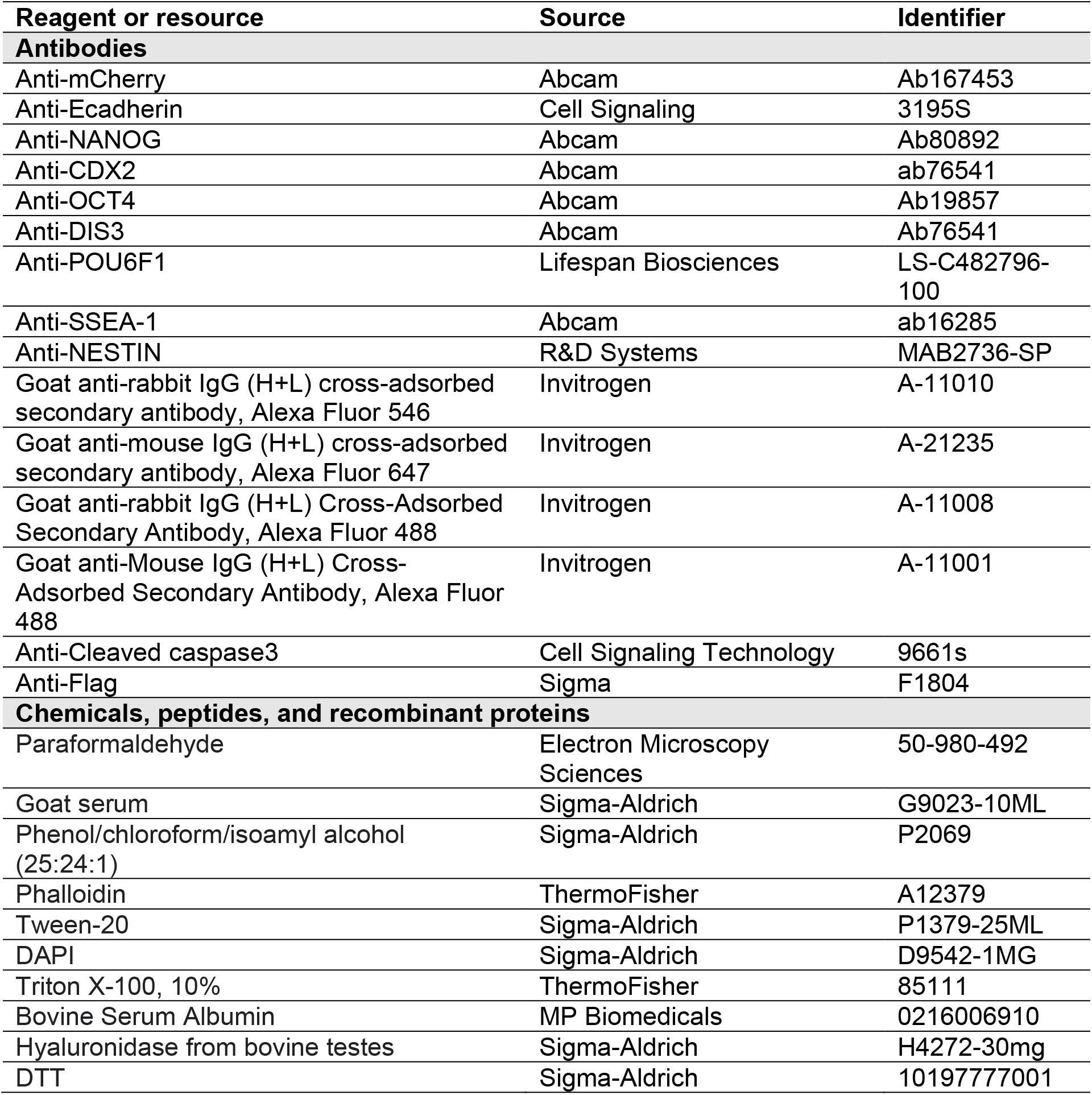
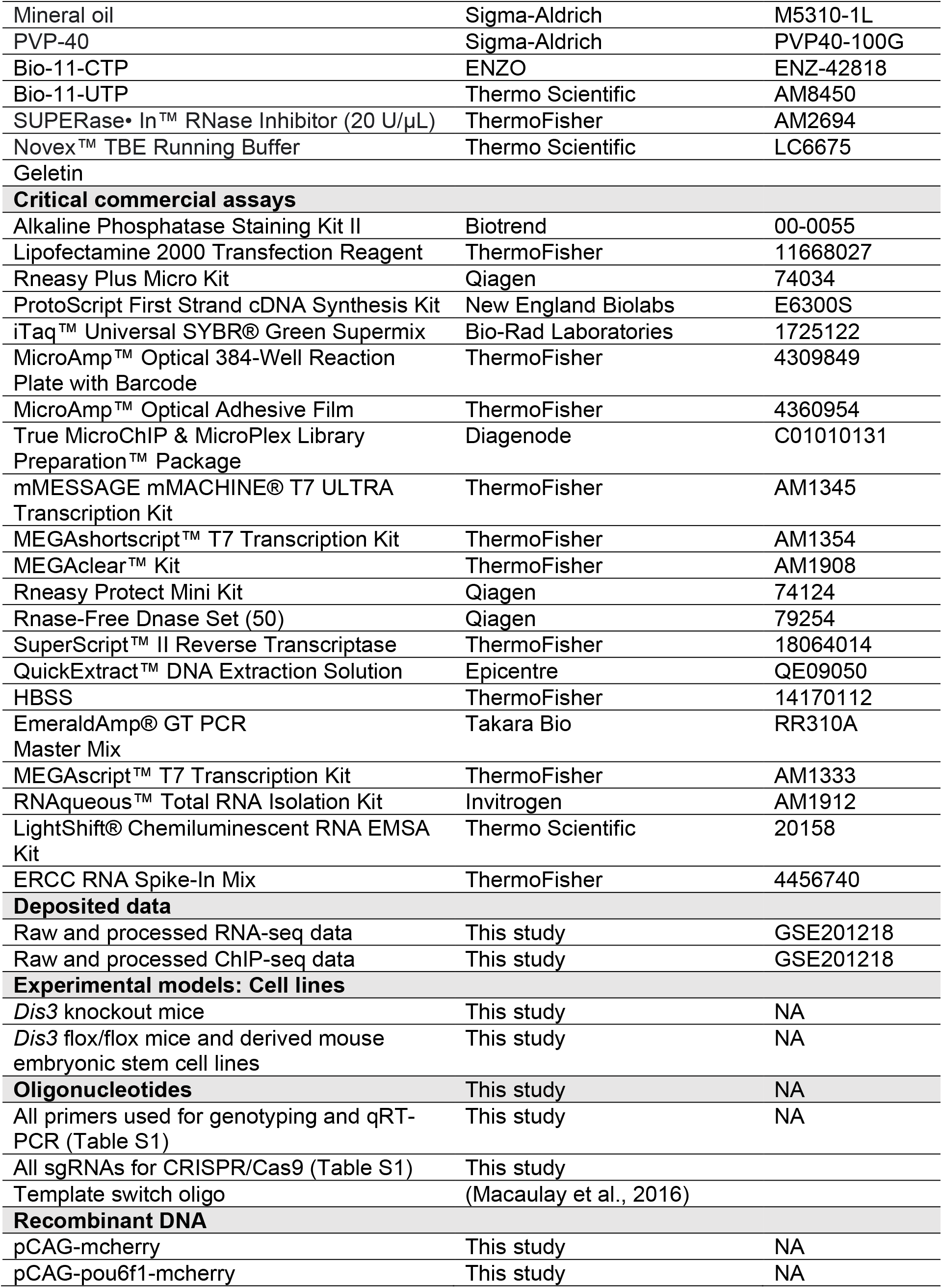
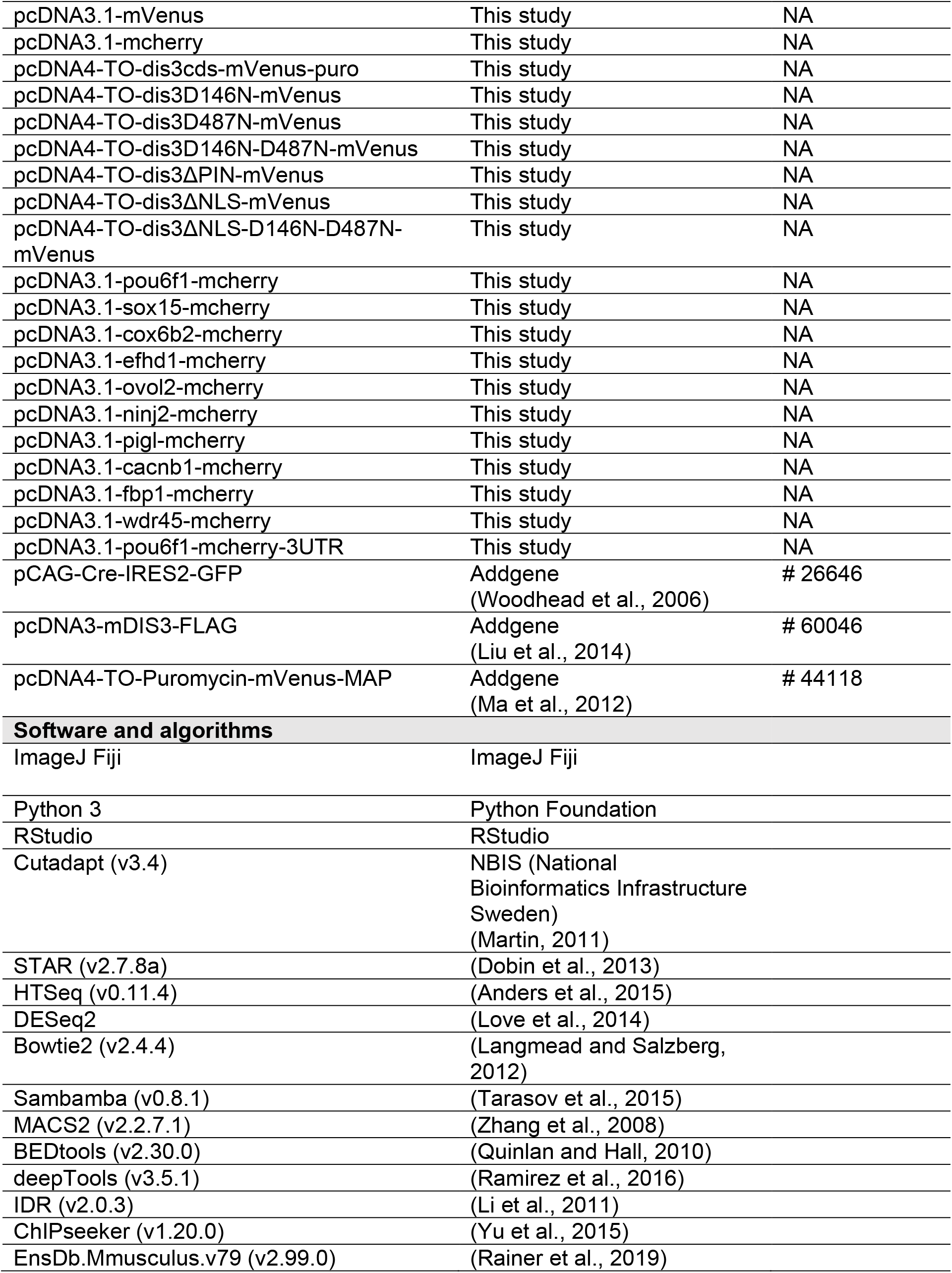
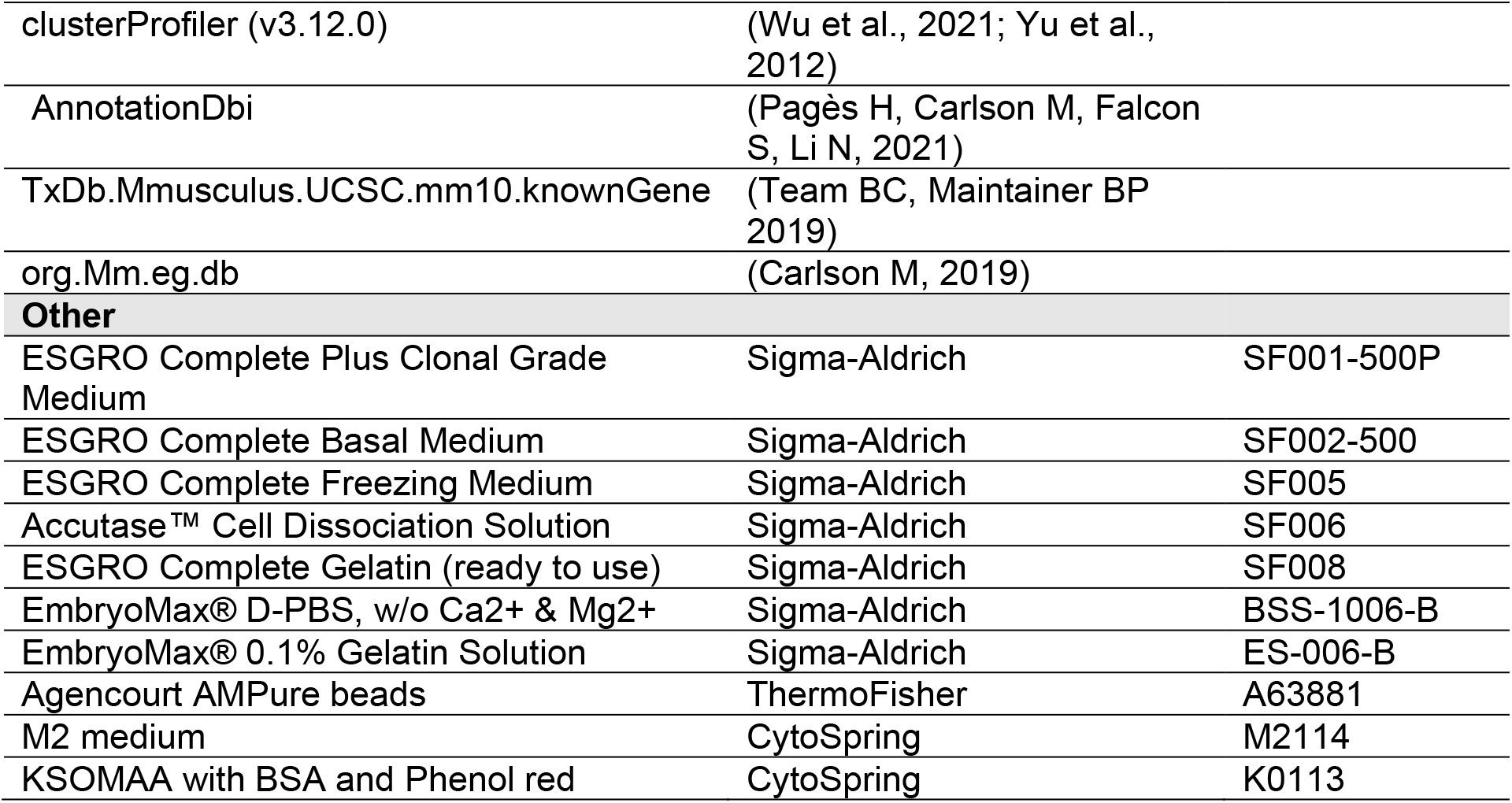

### RESOURCE AVAILABILITY

#### Lead contact

Further information and requests for resources, reagents and data will be addressed by the corresponding authors, Di Wu and Jurrien Dean.

#### Materials availability

All reagents and resources generated in this study are available on request from the lead contacts.

#### Data and code availability

Raw data, processed counts, and peaks of the single embryo RNA-seq the ChIP-seq for POU6F1 in mouse embryonic stem cells were deposited in the NCBI Gene Expression Omnibus (GSE201218). Processed differential analysis results are in supplemental tables. All scripts including R, Jupyter notebooks and Shell were uploaded to https://github.com/Di-aswater. Any additional information, raw images and values are available from the lead contacts upon request.

### EXPERIMENTAL MODEL AND SUBJECT DETAILS

#### Knockout mice generation

*Dis3* knockout mice were generated the same way as the *Dis3* floxed allele (Wu and Dean, 2022). Briefly, two guide RNAs (gRNAs) were designed to delete exons 3-5. The edited gene was repaired via nonhomologous end joining (NHEJ). The gRNAs (50 ng/μl) and *Cas9* mRNA (100 ng/μl) were mixed and microinjected into 1-cell mouse embryos. During microinjection, 1-cell embryos were collected from super ovulated *B6D2_F1_* female mice mated to *B6D2_F2_* stud male mice into M2 medium (CytoSpring M2114). Embryos were treated with hyaluronidase (0.5 mg/ml, Sigma H4272) for 1 min at 30 °C to remove cumulus cells. Injected embryos were washed 10 times with KSOM (CytoSpring K0113) and cultured in KSOM at 37 °C with 5% CO_2_ overnight. 2-cell embryos were transferred into the oviducts of pseudopregnant ICR female mice at 0.5 days post coitus. Two *Dis3* knockout mouse lines were established with different DNA sequences between exon 2 and 6 after deleting exons 3-5 due to imprecise repair.

Genotyping of the mice was performed using EmeraldAmp® GT PCR Master Mix (Takara Bio RR310A). Oligos and genotyping primers are listed in Table S1.

#### *Ex vivo* culture of mouse embryos

For all *ex vivo* culture experiments, female mice at 8-12 wk old were stimulated with 5 IU of PMSG for 48 hr and 5 IU of hCG for 14 hr and mated with male mice. Embryos were dissected in M2 medium, treated with hyaluronidase (0.5 mg/ml) to remove cumulus cells. Embryos were washed 10 times with KSOM and cultured in KSOM at 37 °C with 5% CO_2_ until the desired developmental stage.

#### mESC derivation and directed differentiation

*Dis3^Flox/Flox^* female mice (Wu and Dean, 2022) were stimulated sequentially with PMCG and hCG and mated to *Dis3^Flox/Flox^* male mice. Embryos were flushed from the uterus 3.5 days post coitus with pre-warmed ESGRO Complete Basal Medium (Sigma SF002). All wells of a 24-well plate were treated with 0.5 ml Gelatin solution (Sigma-Aldrich SF008) for 30 min at room temperature. After removing Gelatin, 0.5 ml pre-warmed ESGRO Complete Plus Clonal Grade Medium (Sigma SF001) was added into each well of the plate. Embryos at the blastocyst stage were transferred individually into each well. Embryos were cultured in ESGRO Complete Plus Clonal Grade Medium for 5 days at 37 °C with 5% CO_2_ and outgrowth of cells were observed from some embryos. Cells were treated with Accutase at 37 °C for 1 min to dissociate into single cells. 1-5 single cells were transferred into new ESGRO Complete Plus Clonal Grade Medium, cultured and maintained as stable cell lines. Cells were split when reaching 50% confluence and forming spheroids of 20-50 cells. For differentiation, cells were washed with SF002 medium and cultured in SF002 medium at 37 °C with 5% CO_2_ for 3 days. Transfection of mESCs is described in METHOD DETAILS below.

### METHOD DETAILS

#### Microinjection

The microinjection system consisted of a Zeiss inverted microscope, an Eppendorf Femtojet 4i Injector, a pair of Eppendorf Transferman NK2 micro manipulators. The injector was set to auto mode, the injection time was 0.1 sec, the compensation pressure was 15 and the injection pressure was 500. Needles were pulled from borosilicate glass with filament (BF150-75-10) using a Sutter Flaming/Brown Micropipette Puller. Injection solutions (2 μl) were loaded into the needle with Eppendorf loading pipet Microloader (930001007) from the back. The holding pipets were Eppendorf VacuTip I (5195000036). Embryos were stored in pre-warmed M2 medium on a slide prior to and during microinjection.

#### Plasmid construction

The coding region of mouse *Dis3* gene was amplified from pcDNA3-mDIS3-FLAG, which was a gift from Dr. Zissimos Mourelatos (Addgene plasmid # 60046; http://n2t.net/addgene:60046; RRID: Addgene_60046) (Liu et al., 2014). The T7 promoter sequence and the *Dis3* coding region was cloned into the vector pcDNA4-TO-Puromycin-mVenus-MAP, which was a gift from Dr. Dannel McCollum (Addgene plasmid # 44118; http://n2t.net/addgene:44118; RRID: Addgene_44118) (Ma et al., 2012). The mutant forms of the *Dis3* gene were cloned using Gibson assembly method.

The coding regions of selected DIS3 substrate genes, including *Pou6f1, Sox15, Efhd1, Pigl, Wdr45, Ovol2, Fbp1, Cacnb1, Cox6b2* and *Ninj2* were synthesized as dsDNA using gBlocks Gene Fragments (Integrated DNA Technologies, Inc). The synthesized fragments were cloned into pcDNA3.1-mCherry using Gibson assembly and their sequences are in Table S1.

#### Synthesis of cRNA

For cRNA synthesis from pcDNA4-TO-Puromycin-mVenus-MAP or pcDNA3.1 based plasmids, XbaI (NEB) was used to linearize the plasmid and DNA templates were purified with QIAquick PCR Purification Kit. RNA was synthesized with a mMESSAGE mMACHINE™ T7 ULTRA Transcription Kit (Thermo Fisher AM1345). In each 20 μl reaction, there were 2 μl of 10X T7 Reaction Buffer, 10 μl of 2X NTP/ARCA buffer, 2 μl of T7 enzyme, and 500 ng of linearized DNA template in nuclease free water. Reactions were mixed, incubated at 37 °C for 2 hr and treated with 2 μl of TURBO DNase. Reactions were polyadenylated per manufacturer’s guide. In each polyadenylation reaction, there were 20 μl of 5X E-PAP buffer, 10 μl of 25 mM MnCl2, 10 μl of ATP solution, 36 μl nuclease free water and 20 μl of the *in vitro* transcription reaction mix. Reactions were cultured at 37 °C for 45 min. Synthesized RNAs were purified with a MEGAclear Transcription Clean-Up Kit (Thermo Fisher AM1908).

#### Immunofluorescence staining and confocal microscopy of embryos

Embryos at desired stages were fixed (2% paraformaldehyde, 30 min, 37 °C). Embryos were washed twice with PBVT (PBS-3 mg/ml, polyvinylpyrrolidone-40, 0.1% Tween-20) and incubated with 0.5% Triton X-100 in PBS for 30 min. Embryos were blocked with 5% goat serum in PBVT for 1 hr and incubated in primary antibodies overnight at 4 °C. On the second day, embryos were washed three times with PBVT and incubated with secondary antibodies overnight at 4 °C. On the third day, embryos were washed four times with PBVT, treated with Hoechst for 15 min and immediately transferred into 18-well dish (Ibidi 81817) for confocal microscopy. Images were obtained with a Zeiss LSM 780 microscope and processed with ImageJ Fiji software.

#### Single embryo RNA-seq

RNA-seq of single embryos was performed using GT-seq as described previously (Macaulay et al., 2016). Briefly, *Dis3^+/-^* female mice were stimulated with PMSG followed by hCG and mated with *Dis3^+/-^* male mice. Embryos were collected individually into RLT buffer at E2.5 and E2.75. ERCC spike-in (1:100,000, ThermoFisher 4456740) was added to each sample. RNA was separated by oligo-dT conjugated beads, and the remaining genomic DNA was used for genotyping of each embryo. In total, there were 6 wildtype, 2 heterozygous and 4 homozygous null embryos at the E2.5 stage, and 3 wildtype, 5 heterozygous and 3 homozygous null embryos at the E2.75 stage. RNA was reverse transcribed through template switch method and Superscript II reverse transcriptase (ThermoFisher 18064014). cDNA was amplified for 18 cycles and purified with Agencourt AMPure beads (A63881). Purified cDNA was evaluated with Agilent Bioanalyzer and constructed into sequencing libraries using the Nextera XT Sample Preparation Kit. RNA sequencing was performed at the NIDDK Genomic Core using HiSeq 2500.

#### RNA-seq analysis

RNA-seq raw reads were quality trimmed using Cutadapt (v3.4) with “-m 10 -j 8 -q 20,20 parameters” (Martin, 2011). Reads were mapped to GRCm38 using STAR (v2.7.8a) with default parameters to generate sorted BAM files (Dobin et al., 2013). BAM files were counted by HTSeq (v0.11.4) with “-m intersection-strict -f bam -s no” parameters (Anders et al., 2015). Count files were used for differential analysis with DESeq2 to obtain differentially expressed genes (Love et al., 2014). Differential analysis was performed individually at each stage and two stages were merged by genotype to perform differential analysis between genotypes. Volcano plots were generated with Jupyter-notebook. All scripts were posted to Github (https://github.com/Di-aswater).

#### Transfection of mESCs

Derived mESC transfection was performed with Lipofectamine 2000 Transfection Reagent (ThermoFisher 11668027). Cells were treated with Accutase at 37 °C for 1 min to dissociate into single cells. Cells were washed by ESGRO Complete PLUS Clonal Grade Medium and ready for transfection (500 μl for a well of a 24-well plate).

For a well of a 24-well plate, DNA (0.5 μg) was diluted into 50 μl of ESGRO Complete PLUS Clonal Grade Medium. Lipofectamine 2000 (1 μl) was diluted into 50 μl of ESGRO Complete PLUS Clonal Grade Medium and incubated for 5 min at room temperature. The diluted DNA and the diluted Lipofectamine 2000 were mixed and incubated for 5 min at room temperature. Transfection mixture was added to cells and mixed thoroughly by rotating up and down for 10 times. Cells were then cultured at 37 °C for 30 min. Cells were centrifuged at 1300 rpm for 5 min, dispensed into new ESGRO Complete PLUS Clonal Grade Medium.

#### Immunofluorescence staining and confocal microscopy of mESCs

Cells were fixed using 4% paraformaldehyde (30 min, room temperature). Cells were washed twice with PBS and permeabilized with 0.5% Triton X-100 for 30 min. Cells were blocked with 5% goat serum diluted in PBS-0.1% Tween 20 for 1 hr. Cells were incubated with primary antibodies overnight at 4 °C. On the second day, cells were washed with PBS-0.1% Tween 20 for 20 min X4 and incubated in secondary antibodies. On the third day, cells were washed with PBS-0.1% Tween 20 for 20 min X4, incubated with DAPI and imaged using the Zeiss LSM 780 confocal microscope. At each step, cells were collected by centrifuging at 1300 rpm for 5 min.

#### Quantitative RT-PCR of mESCs

Cells were washed with PBS and collected into RLT buffer. Total RNA was extracted with a Qiagen RNeasy Plus Micro Kit (74034). Total RNA was used for reverse transcription with a ProtoScript First Strand cDNA Synthesis Kit (New England Biolabs E6300S) in a 20 μl reaction. In each reaction, there were 1 μg of total RNA, 2 μl of d(T)23VN oligos (50 μM) in nuclease free water (8 μl in total). The mixture was denatured for 5 minutes at 70°C, spined down briefly and put on ice. In each reverse transcription reaction, there were 10 μl of AMV Reaction Mix, 2 μl of AMV Enzyme Mix, and 8 μl of RNA mix prepared above. Reactions were incubated at 25°C for 5 min, 42°C for 1.5 hr and 80°C for 5 min. The generated cDNA was diluted to 50 μl by adding 30 μl of nuclease free water. qRT-PCR was performed with iTaq™ Universal SYBR® Green Supermix (Bio-Rad Laboratories 1725122). 10 μl reactions were performed in a MicroAmp™ Optical 384-Well Reaction Plate with Barcode (4309849). In each reaction, there were 5 μl of 2X Supermix, 0.25 μl of each primer (10 μM), 0.5 μl of DNA (1-10 ng/μl) and 4 μl water. qPCR reactions were performed using QuantStudio 6 Flex Real-Time PCR System (Thermo Fisher Scientific).

#### ChIP-seq

mESCs were transfected with pCAG-mCherry or pCAG-pou6f1-mcherry. Each transfection sample was performed using 15 μg of DNA and 30 μl Lipofectamine 2000 (details described above) in a T75 cell culture flask of about 1,000,000 cells. Each sample had three biological replicates. Cells were dissociated by Accutase and collected into lobind tubes 48 hr after transfection. ChIP-seq libraries were prepared using True MicroChIP & MicroPlex Library Preparation™ Package (C01010131) according to the manufacturer’s instructions. Briefly, cells were washed with PBS and fixed by 1% formaldehyde (15 min, room temperature). Crosslink was stopped by adding glycine (115 μl per 1 ml reaction) and incubated for 5 min at room temperature. Cells were washed twice with HBSS (ThermoFisher 14170112), collected, and for frozen overnight at −80 °C. On the second day, cells were lysed and sonicated using the Bioruptor for 10 cycles. Sheared DNA was purified using phenol/chloroform/isoamyl alcohol (25:24:1) (Sigma-Aldrich P2069). 20 μl from each sample was removed as the input sample and held at −80 °C. 1 μg of the anti-mCherry antibody or rabbit IgG was added into each sample. Samples were incubated overnight at 4 °C. On the next day, magnetic beads were pre-treated and added into each sample and incubated for 2 hr at 4 °C. Samples were washed, de-crosslinked and purified using phenol/chloroform/isoamyl alcohol (25:24:1). Quality of the samples was evaluated by Bioanalyzer 2100. Samples were used for library preparation following the MicroPlex Library Preparation manual, purified with Agencourt AMPure XP beads and pooled for sequencing. The sequencing was performed at the NIDDK Genomics Core using HiSeq 2500.

#### ChIP-seq analysis

Original ChIP-seq fastq files were aligned to GRCm38 using Bowtie2 (v2.4.4) (Langmead and Salzberg, 2012). Generated SAM files were converted to BAM files, sorted, and filtered for uniquely mapping reads using Sambamba (v0.8.1)(Tarasov et al., 2015). Peaks were called using MACS2 (v2.2.7.1) (Zhang et al., 2008), and the anti-IgG sample and anti-mCherry of mCherry transfected samples used as controls, respectively. Each biological replicate was processed individually to obtain peaks. Consensus peaks were generated using BEDtools (v2.30.0) (Quinlan and Hall, 2010). When plotting heatmaps, coverage values (CPM normalized) were calculated from BAM files using deepTools (v3.5.1) (Ramirez et al., 2016), and processed with computeMatrix and plotHeatmap functions of deepTools.

#### Electrophoretic Mobility Shift Assay (EMSA)

*Pou6f1* probe was synthesized by *in vitro* transcription. The *Pou6f1* exon 8 was selected as the probe because of the DIS3 occupancy in the PAR-CLIP seq result (Szczepinska et al., 2015). The probe is 121 nt long (Table S1), and its template was synthesized as ssDNA by adding T7 in the 5’ end and T3 in the 3’ end (Table S1). The ssDNA was PCR amplified by T7 and T3 primers. Purified dsDNA was used as the template for *in vitro* transcription using MEGAscript™ T7 Transcription Kit (ThermoFisher AM1333). For unlabeled probe synthesis, each reaction had 2 μl of 75 mM ATP, GTP, CTP, UTP, respectively, plus other essential components per manufacture’s user guide. For Biotin-labeled probe synthesis, each reaction had 2 μl of 75 mM ATP, 2 μl of 75 mM GTP, 1.5 μl of 75 mM CTP, 1.5 μl of 75 mM UTP, 3.75 μl of 10 mM Bio-11-CTP (ENZO ENZ-42818), 3.75 μl of 10 mM Bio-11-UTP (Thermo Scientific AM8450) plus other essential components. Synthesized probes were purified by Ambion RNAqueous™ columns (Invitrogen AM1912).

To prepare the crude cell extract, pcDNA4-TO-dis3ΔNLS-D146N-D487N-mVenus protein was transfected into the derived mESCs and cultured for 48 hr. Cells were collected, washed and transferred to newly prepared lysis buffer (150 mM NaCl, 1% Triton X-100, 50 mM Tris-Cl, Protease inhibitor cocktail, pH 7.4). Cell-lysis solution was mixed by vortex and incubated on ice for 30 min. Lysis was frozen at −80C overnight, thawed, diluted to 1 mg/ml, and centrifuged at 12,000 rpm for 10 min to remove debris before use. EMSA was performed using LightShift® Chemiluminescent RNA EMSA Kit (Thermo Scientific 20158). DNA Retardation Gels (6%) was used (Thermo Scientific EC6365BOX). In each 20 μl reaction, there were 1 μl of RNase inhibitor (ThermoFisher), 2 μl of 10X binding buffer, 1 μl of 50% glycerol, 1 μl of 100 mM MgCl2, 1 μl of 1 ng/μl bio-probe, 2 μl of 0.1 ng/μl unlabeled probe (when necessary), 1 μl of protein extract and 1 μl of anti-Flag (Sigma F1804) in ultrapure water. Probes were denatured by incubation at 80 °C for 5 min, spined down and put on ice right before use. Reactions were incubated at room temperature for 30 minutes. 5 μl of 5X loading buffer was added to each 20 μl reaction and mixed by pipetting. 20 μl of reaction was loaded onto each well for electrophoresis through 6% non-denaturing polyacrylamide gel in 0.5X TBE buffer, transferred to a nylon membrane (Thermo Scientific AM10100), crosslinked to the membrane by a commercial UV-light crosslinking instrument (120 mJ/cm^2^ for 60 s). Membrane was incubated by the blocking buffer, washing buffer, substrate equilibration buffer and detected per user’s guide.

### QUANTIFICATION AND STATISTICAL ANALYSIS

Statistical analyses in Figure 1-3 were performed using two-tailed Student’s t-test. P values are labeled in each figure and the figure legends. All group photos were from one assay out of at least three repeated assays. All single embryo photos were the most representative picture from all repeated experiments. The number of samples/embryos is labeled in each figure or figure legend. In the RNA-seq analysis, P-adj <0.1 and log2 fold change >1 was implemented to filter the significantly changed genes.

## ACKNOWLEDGMENTS

We thank the critical reading of the manuscript by all the members of the Jurrien Dean lab. We appreciate the bioinformatic training and discussion from Cameron Palmer, Ph.D.

## AUTHOR CONTRIBUTIONS

D.W. and J.D. conceived the project. D.W. designed and performed the experiments and bioinformatic analysis. D.W. and J.D. wrote manuscript.

## SUPPLEMENTAL INFORMATION

**Figure S1.**
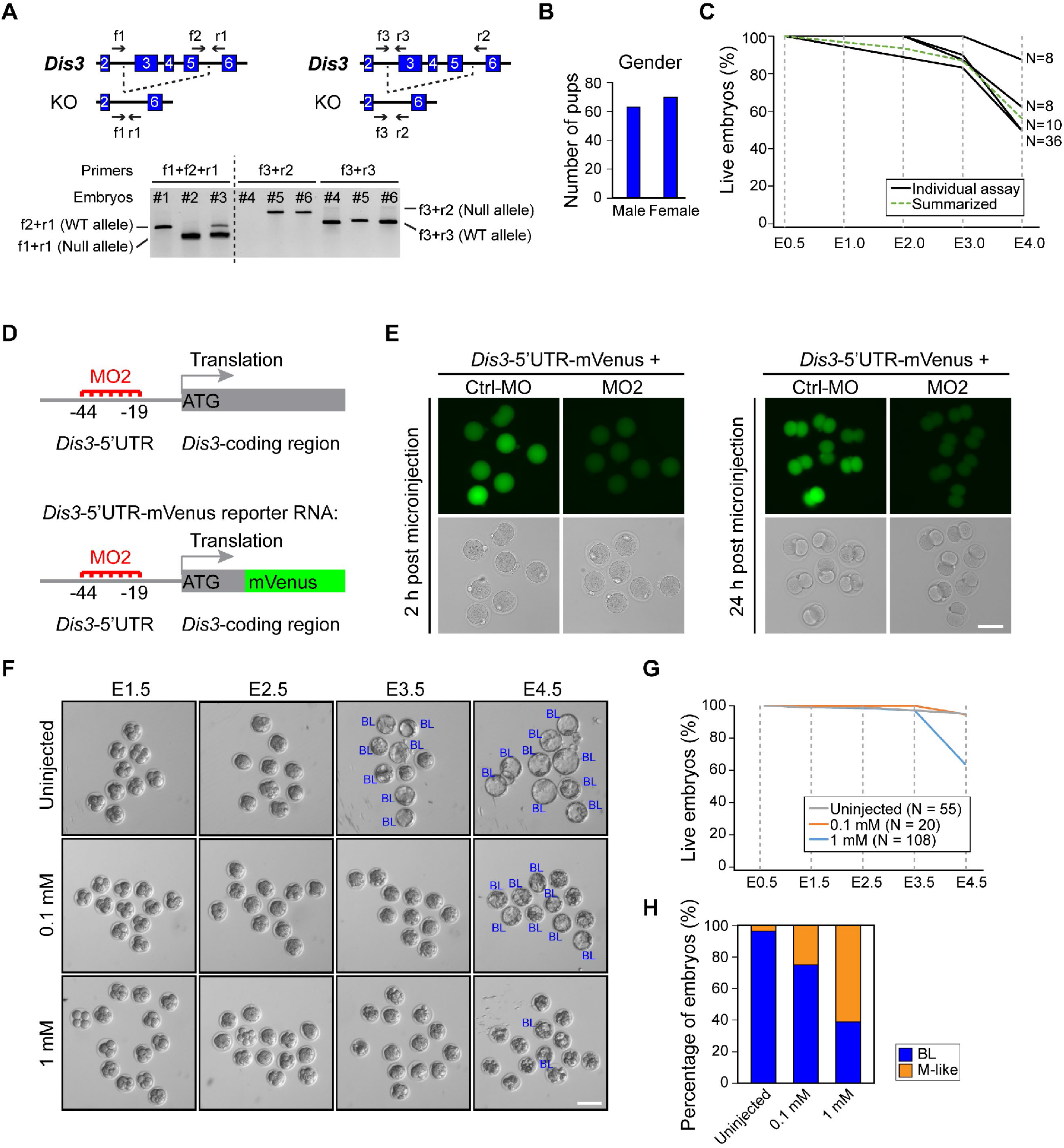
*Dis3* Null Embryos Arrest at the Morula Stage. (A) Schematic and DNA electrophoresis results of genotyping. Two genotyping strategies are illustrated. Arrows label the positions of primers used. In the first strategy (left), f2+r1 is specific to the WT allele and f1+r1 is specific to the null allele. In the second strategy (right), f3+r2 is specific to the WT allele and f3+r2 is specific to the Null allele. #1-6: six embryos. #1 and #4: WT embryos; #2: null embryo; #3, #5 and #6: het embryos. (B) Bar graph showing gender distribution of the pups born from *Dis3* heterozygous mating. (C) Line graph showing the viability of embryos derived from *Dis3* heterozygous mating in pre-implantation development. Black lines: four assays; green dash line: summary of the four assays. (D) Schematic of *Dis3* MO2 in blocking *Dis3* mRNA translation (top) and schematic of the reporter RNA to test MO2 efficiency (bottom). (E) Fluorescent and bright field imaging of wildtype embryos injected with reporter RNA and *Dis3* MO2 and control MO (Ctrl-MO). (F) Bright field live imaging of wildtype embryos microinjected with *Dis3* MO2. (G and H) Line graph and bar graph showing the survival rate of *Dis3* MO2-injected embryos and the stage of embryos at E3.5. Scale bars: 100 μm in E, F.

**Figure S2.**
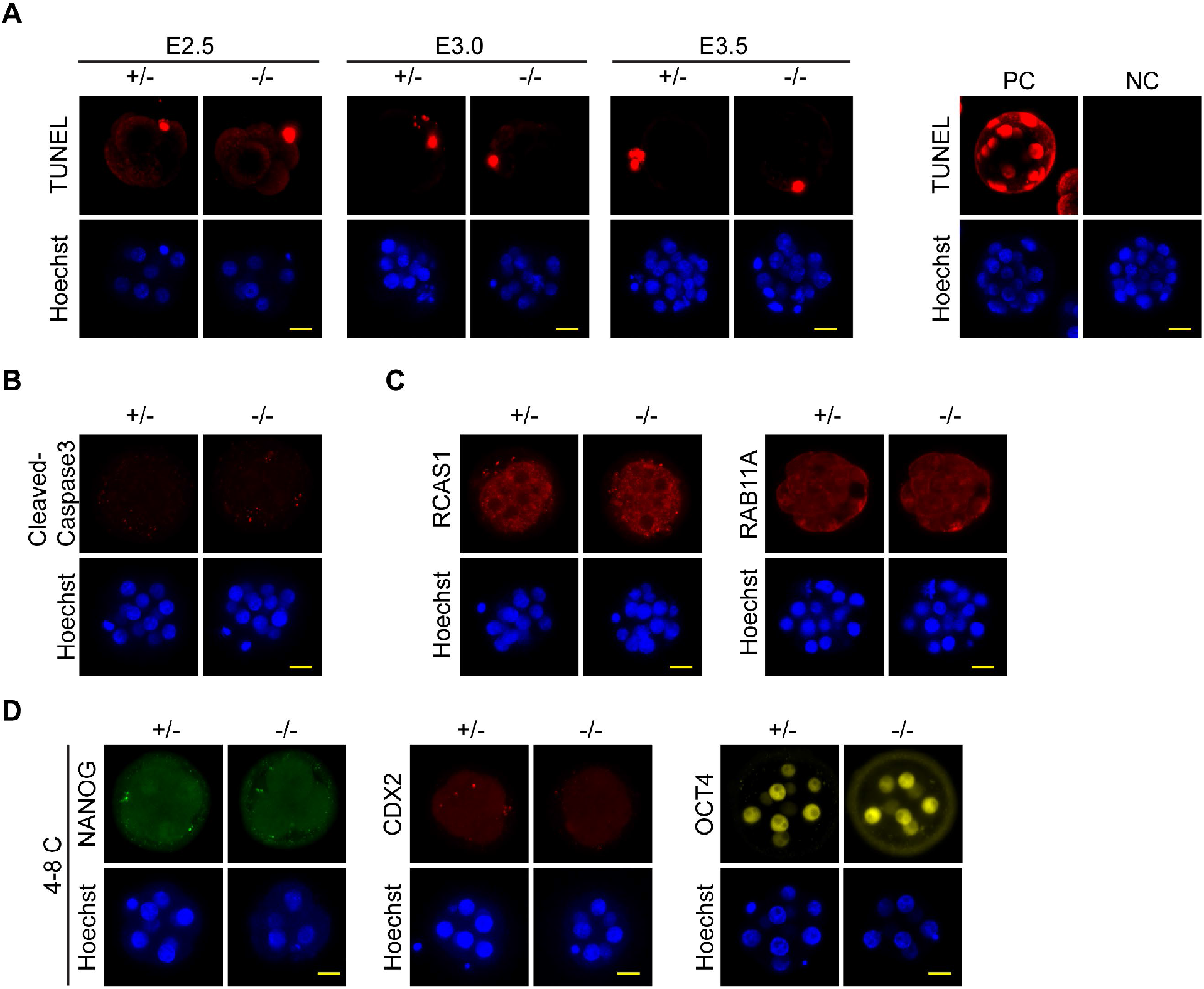
Immunostaining of *Dis3* Null Embryos for Apoptosis, Endomembrane Integrity and Differentiation. (A) Immunofluorescent TUNEL assay of *Dis3* het (+/-) and null (-/-) embryos. PC: positive control of embryos treated with DNase I (3 U/μL). NC: negative control of embryos without staining. (B and C) Immunofluorescence staining of cleaved caspase 3, RCAS1 and RAB11A. (D) Immunofluorescence of NANOG, CDX2 and OCT4 in *Dis3* het and null embryos at the 4-8 cell stage. Scale bars: 20 μm.

**Figure S3.**
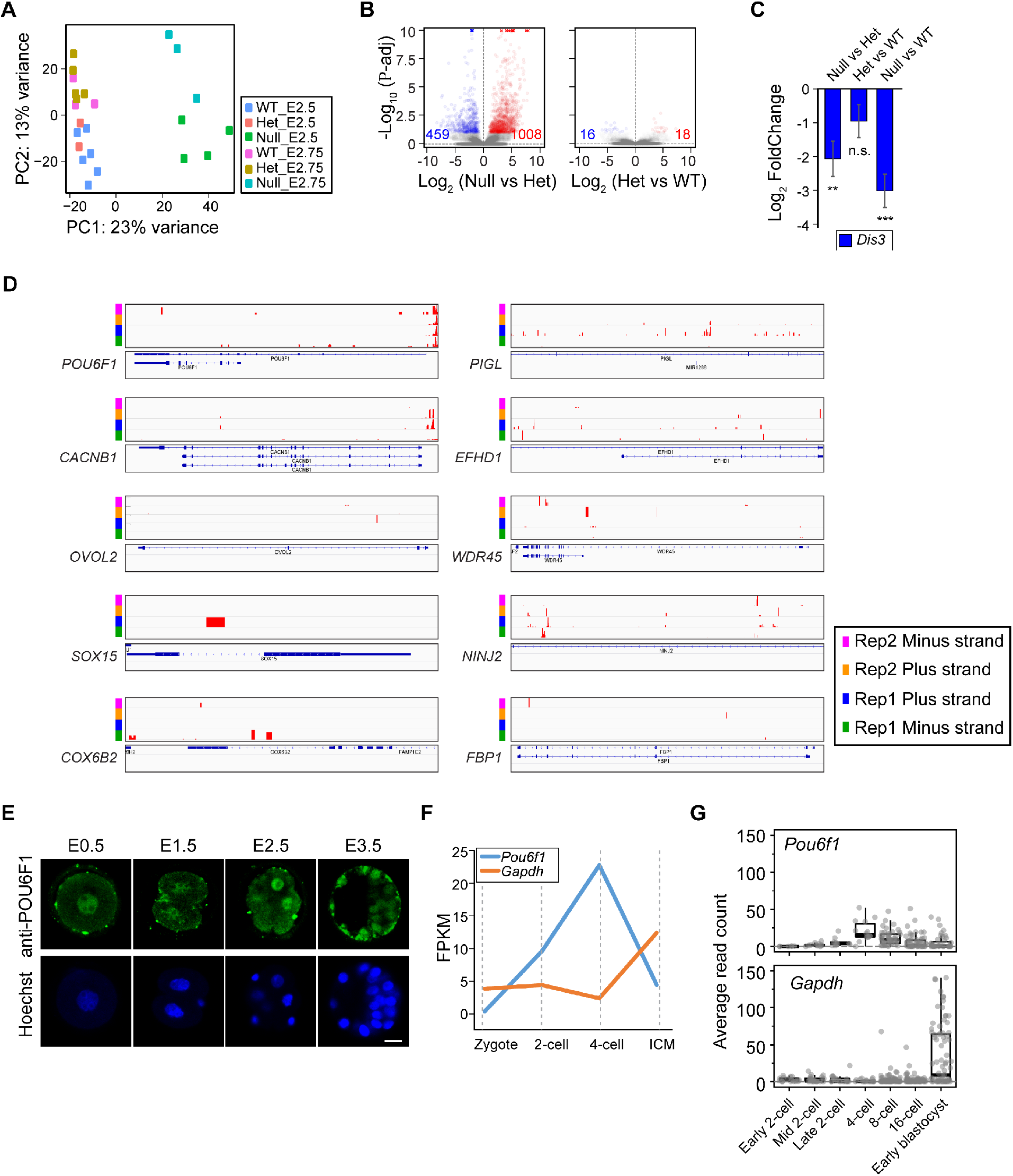
Identification of *Pou6f1* from Single Embryo RNA-seq. (A) Dot plot of principle component analysis (PCA) of single embryo RNA-seq samples. (B) Volcano plots of differentially expressed promoter upstream transcript (PROMPTs) in single embryo RNA-seq of combined E2.5 and E2.75 stages. (C) Bar graph of changes in abundance of *Dis3* transcript in RNA-seq. ** P-adj < 0.01, *** P-adj < 0.001, n.s., not significant. (D) Integrated genomic viewer of PAR CLIP-seq results of the 10 substrate genes (Szczepinska et al., 2015). Reference used is the human genome. Four tracks shown include two replicates of plus strand and minus strand. (E) Immunofluorescence staining of POU6F1 in wildtype embryos from E0.5 to E3.5. (F) FPKM (fragments per kilobase million) of *Pou6f1* and *Gapdh* from GSE57249 (Biase et al., 2014) RNA-seq results of blastomeres at different stages during mouse pre-implantation development. (G) Average read counts of *Pou6f1* and *Gapdh* from GSE45719 (Deng et al., 2014) single cell RNA-seq from 2-cell stage to blastocyst stage. Scale bar: 20 μm.

**Figure S4.**
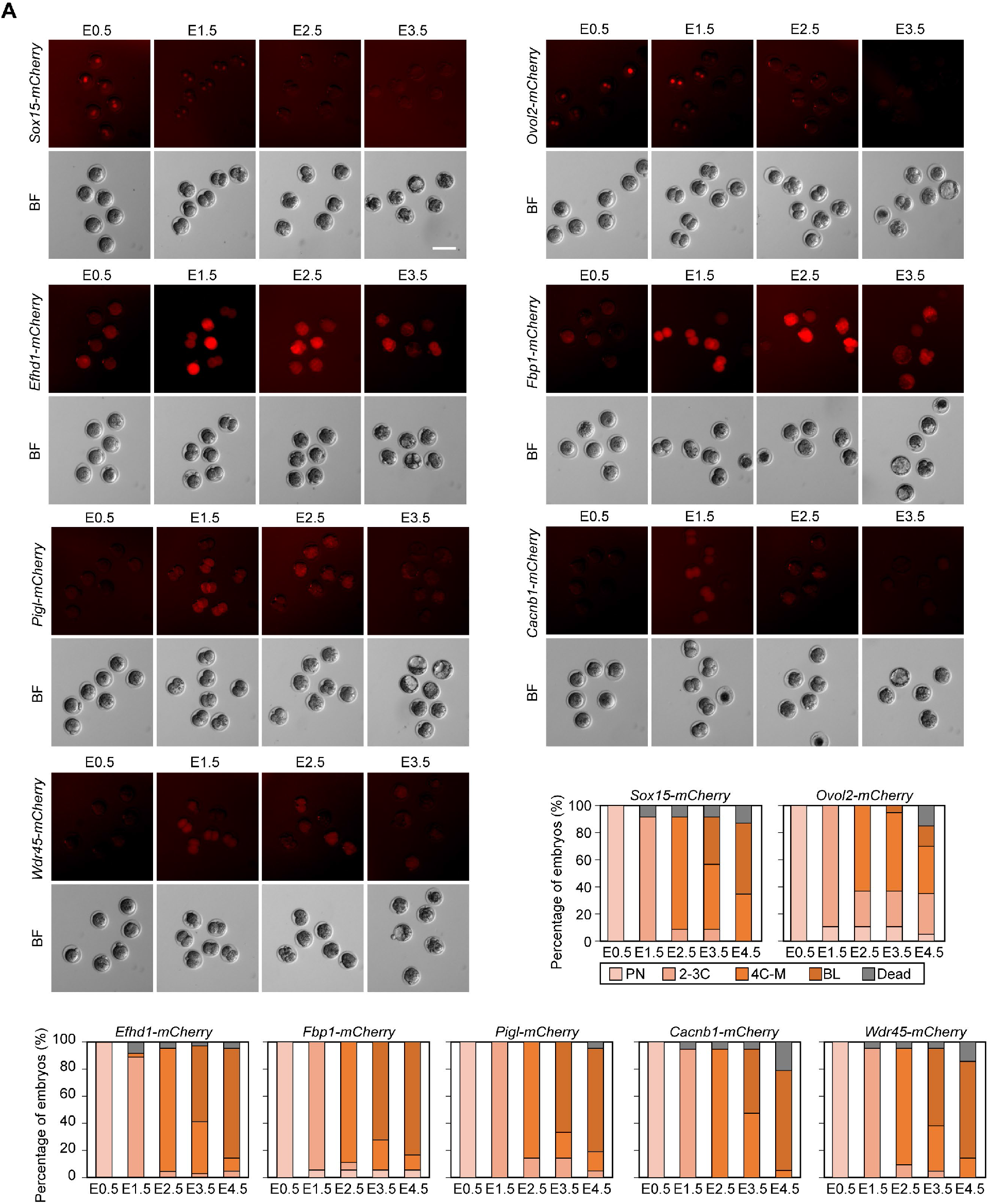
Overexpression of Selected DIS3 Substrates Does Not Phenocopy *Dis3* Null Embryos. (A) Fluorescence and bright field imaging of wildtype embryos injected with potential DIS3 substrates individually: *Sox15-mCherry* (N=23), *Ovol2-mCherry* (N=26), *Efhd1-mCherry* (N=22), *Fbp1-mCherry* (N=22), *Pigl-mCherry* (N=21), *Cacnb1-mCherry* (N=19), *Wdr45-mCherry* (N=21). Bar graphs show the stages of embryos at different time points. Scale bar: 100 μm.

**Figure S5.**
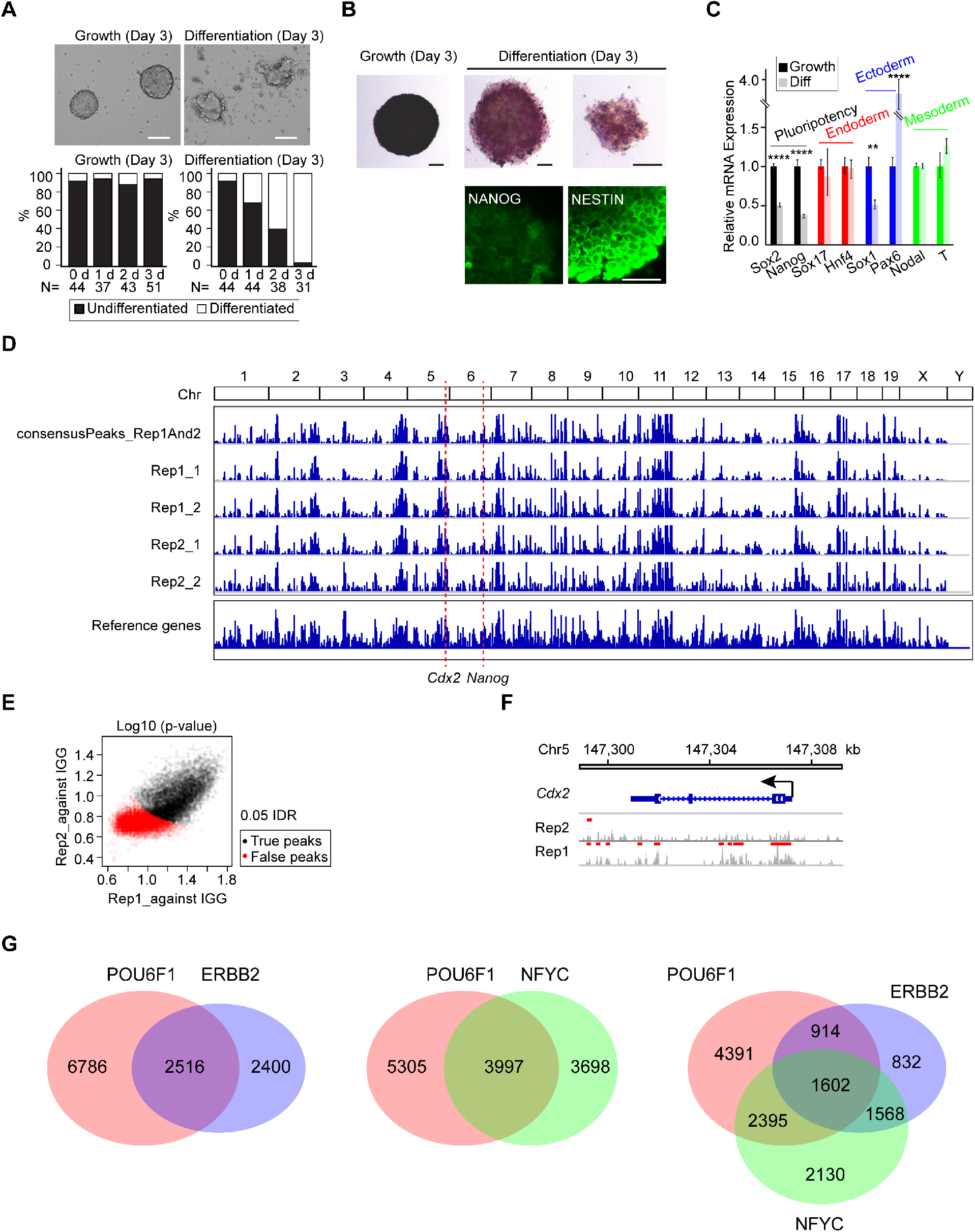
Identification of POU6F1 Occupied Genes Using ChIP-seq of mESC. (A) Bright field images of derived mESCs cultured in growth condition (self-renewal factors available) and differentiation condition (basal medium, without self-renewal factors). Bar graph shows the percentage of colonies counted in undifferentiated or differentiated morphology. Scale bars: 100 μm. (B) Differentiation of mESC labeled by AP staining and immunofluorescence staining of NANOG and NESTIN. Scale bar: 50 μm. (C) Bar plot of quantitative RT-PCR of different marker genes in growth and differentiation (Diff) conditions of derived mESCs. **** Sox2, Nanog, Pax6, P< 0.001; ** Sox1, P< 0.01; two-tailed Student’s t-test. (D) Integrated genomic viewer of ChIP-seq results of two biological replicates. For each replicate (Rep1 and Rep2), peaks are called either against IgG of the *Pou6f1-mCherry* transfected cells (Rep1_1, Rep2_1) or against mCherry of the *mCherry* transfected (Rep1_2, Rep2_2). (E) Dot plot showing IDR analysis results. Each dot is a peak from two replicates (Rep1 and Rep2) called against IGG. Black dots are IDR value less than 0.05, which are suggested as true reproducible peaks; red dots are IDR values larger than 0.05, which are suggested as false reproducible peaks. (F) POU6F1 peaks on *Cdx2*. Red bars: peaks called against IGG samples. Venn diagrams showing the number of overlapped genes between POU6F1 and ERBB2 (left), POU6F1 and NFYC (middle), POU6F1, ERBB2 and NFYC (right).

